# SurfStamp: 3D Printer Compatible Molecular Surface Representation

**DOI:** 10.1101/2020.10.29.360701

**Authors:** Toshiyuki Oda

**Affiliations:** Independent software developer

## Abstract

SurfStamp is an application that is used to generate textures for surface models of proteins. The textures contain information about surface residues and the information is drawn directly on the 3D object of the models. This approach is more intuitive than the labeling functions that most three-dimensional (3D) structure viewers use to show residue information. Therefore, the use of this application enables researchers, readers, or audiences to easily determine which residues are contributing the surface they are focusing on.

**Availability:** The application is provided under the open-source Apache License Version 2.0 (http://www.apache.org/licenses/LICENSE-2.0). The application and source code are available from https://github.com/yamule/SurfStamp-public/releases. The application is also available as a plugin for the recent versions of PyMOL (Schrodinger, 2015, https://github.com/schrodinger/pymol-open-source) from https://github.com/yamule/SurfStamp-pymol-plugin.

## Introduction

Since visualization of protein structures in 3D space is one of the fundamental ways to analyze properties and function of proteins, various techniques have been developed and are now in widespread use; e.g., tube models, ball-and-stick models, surface models, and so forth (Gu and Bourne, 2009). Among them, molecular surface model is one of the more important techniques; because the properties of the surface regions determine their binding affinities to other molecules (e.g. small chemical compounds, DNAs or other proteins) and these affinities are strongly related to the functions of proteins.

For this reason, popular 3D structure visualization software products such as PyMOL (Schrodinger, 2015), VMD (Humphrey, et al., 1996), and JMol (http://www.jmol.org/) each have functions that can be used to construct molecular surface models. Using these software products, researchers can see protein shapes and interesting regions such as cavities, hydrophobic regions, or charged clusters, which will expand their knowledge and provide clues for further experiments.

However, when examining surface models, it is often difficult to see which residues contribute to the regions under examination. While most molecular visualization software is capable of adding labels that represent residues, those labels often need manual rearrangement in order to make them understandable when they are shown with a surface.

In this paper, I introduce SurfStamp, a software tool for generating textures which show residue types and numbers on a surface model. Using this application, residue information is drawn directly on the surface object, which makes it easy to understand which residues are contributing to the regions of interest. In addition, since the residue information is included in a 3D object, it can be utilized in 3D printing, which is useful in scenes of research and pedagogy (Da Veiga Beltrame, et al., 2017).

## Implementation

### Work flows

The work which SurfStamp does is quite simple. It can be broken down into 5 or 6 steps as follows.

1. Surface model generation SurfStamp creates models using the similar approach of molecular surface mode of EDTSurf (Xu and Zhang, 2009), however, the integrity of the algorithm has not been tested. Users can import surface models created by other popular software products such as EDTSurf or PyMOL.
2. Grouping faces (polygons) All faces in those geometries are grouped according to their nearest residues. The distances between the center point of each face and any atoms of residues are used.
3. Topology validation The topologies of grouped faces are checked. When the default setting is used, each group is separated into two groups heuristically if there are large areas that would not be seen from one side. If there are loop structures such as tori, which cause failures of UV unwrapping, those structures are removed. Structures such as sharp hills are also removed because they result in extremely skewed textures.
4. UV unwrapping The faces are unwrapped onto a two-dimensional (2D) plane and the texture coordinates are decided.
5. Texture creation The unwrapped faces are then painted according to the coloring scheme, after which text showing the residue types and numbers are placed inside the unwrapped textures. If the “-extrude” option is not present, the process stops here and result files are created.
6. Extrusion With the “-extrude” option (extrusion mode), SurfStamp doesn’t create texture files. It extrudes residue information from surface model as solid objects using the texture data created in step 5.

### Coloring scheme

When the default setting is in place, a coloring scheme similar to ClustalX (Thompson, et al., 2002) is used. The Hydropathy index (Kyte and Doolittle, 1982) and Charge (Zimmerman, et al., 1968) extracted from AAIndex Database (Kawashima, et al., 2008) are provided as presets. The CPK coloring scheme (Corey and Pauling, 1953; Koltun, 1965), B-factor or occupancy columns can be used for atom, too. However, it should be noted that users can apply their own original coloring schemes by using a simple script (the details are written in the readme.txt file provided with the application).

### Input file format

For 3D polygon objects, SurfStamp accepts Wavefront .obj format files (https://en.wikipedia.org/wiki/Wavefront.obj_file), Polygon file (.ply) format files (https://en.wikipedia.org/wiki/PLY_(file_format)), and binary STL format (https://en.wikipedia.org/wiki/STL_(file_format)) as inputs. I optimized the software to import files by PyMOL (v0.99, https://sourceforge.net/projects/pymol/files/Legacy/), which is one of the most popular visualization software tools, and EDTSurf, which is a character user interface (CUI) based molecular surface generation tool that can be used for batch processes. However, it is hoped that the files generated by other software can be processed as well.

For atom coordinates, standard file formats for protein structures, the Protein Data Bank (PDB) file format, and the macromolecular Crystallographic Information File (PDBx/mmCIF) format are supported. For ligand molecules, SDF file format is also supported.

### Output file format

The default mode of SurfStamp produces 3D graphics files in .obj and Material library (.mtl) formats (https://en.wikipedia.org/wiki/Wavefront_.obj_file#Material_template_library), while texture image files are provided in Portable Network Graphics (.png) formats (https://www.w3.org/TR/PNG/). Since these formats are widely used in the 3D or 2D graphics field, they are supported by most 3D or 2D graphics software applications.

In addition, texture images are also provided in the Scalable Vector Graphics (.svg) format (https://www.w3.org/TR/SVG/), which can be processed with vector draw software such as Inkscape (https://inkscape.org/). Using this format (and its related software), users can change the texts, fonts, colors, and resolution rates more easily than can be done with raster-type formats such as .png.

In extrusion mode, it exports objects in binary STL format.

### Additional features

SurfStamp can produce ball-and-stick molecule models. This feature was added to allow researchers to determine whether a molecular surface has been successfully created. (Since some visualization software produce 3D objects according to its camera position, the coordinates of produced objects sometimes differ from coordinates of atoms.) Note, however, in addition to this debugging purpose, the feature is also useful for showing the positions of ligands, as can be seen in Fig. 1.

**Figure 1.**
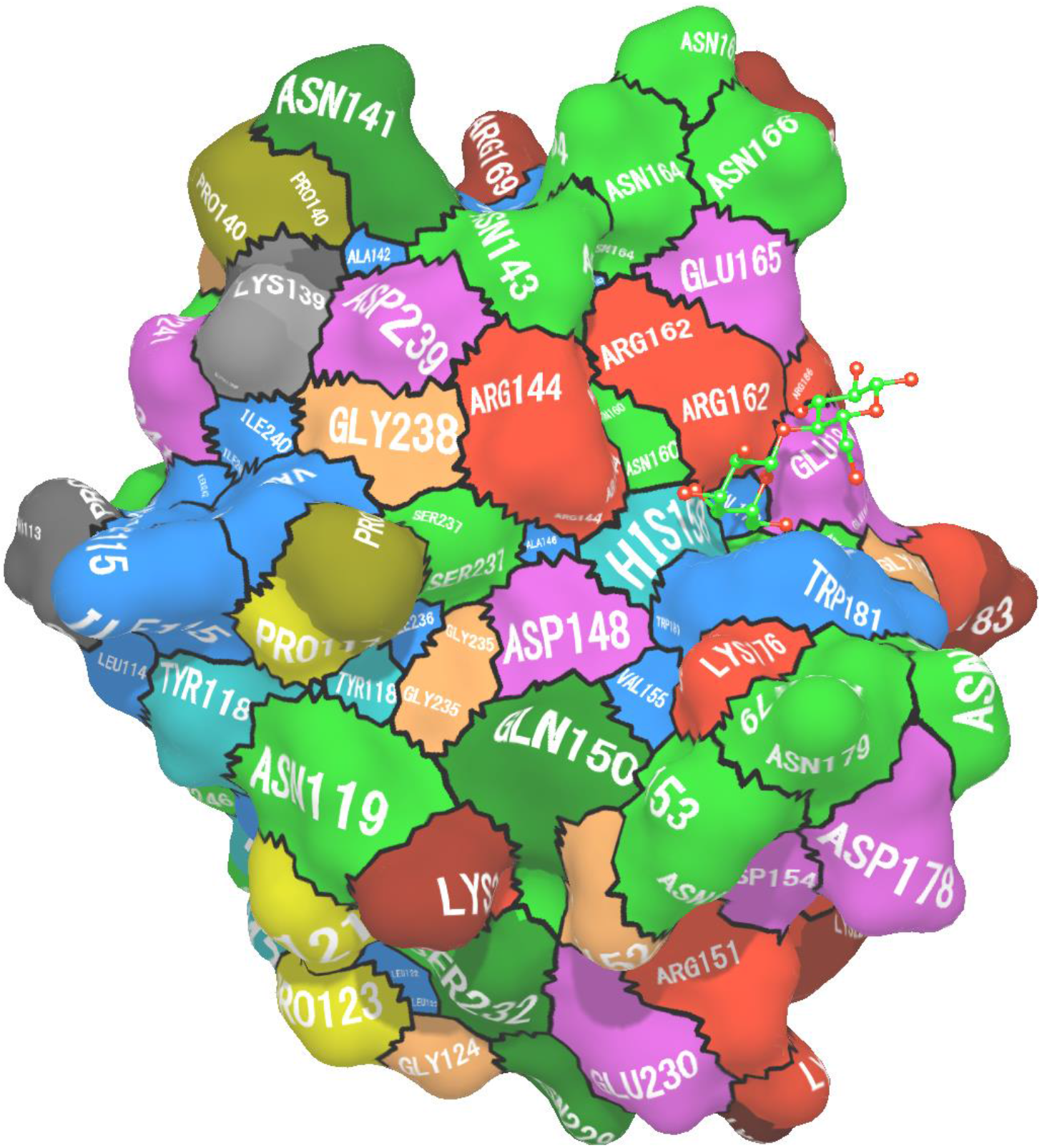
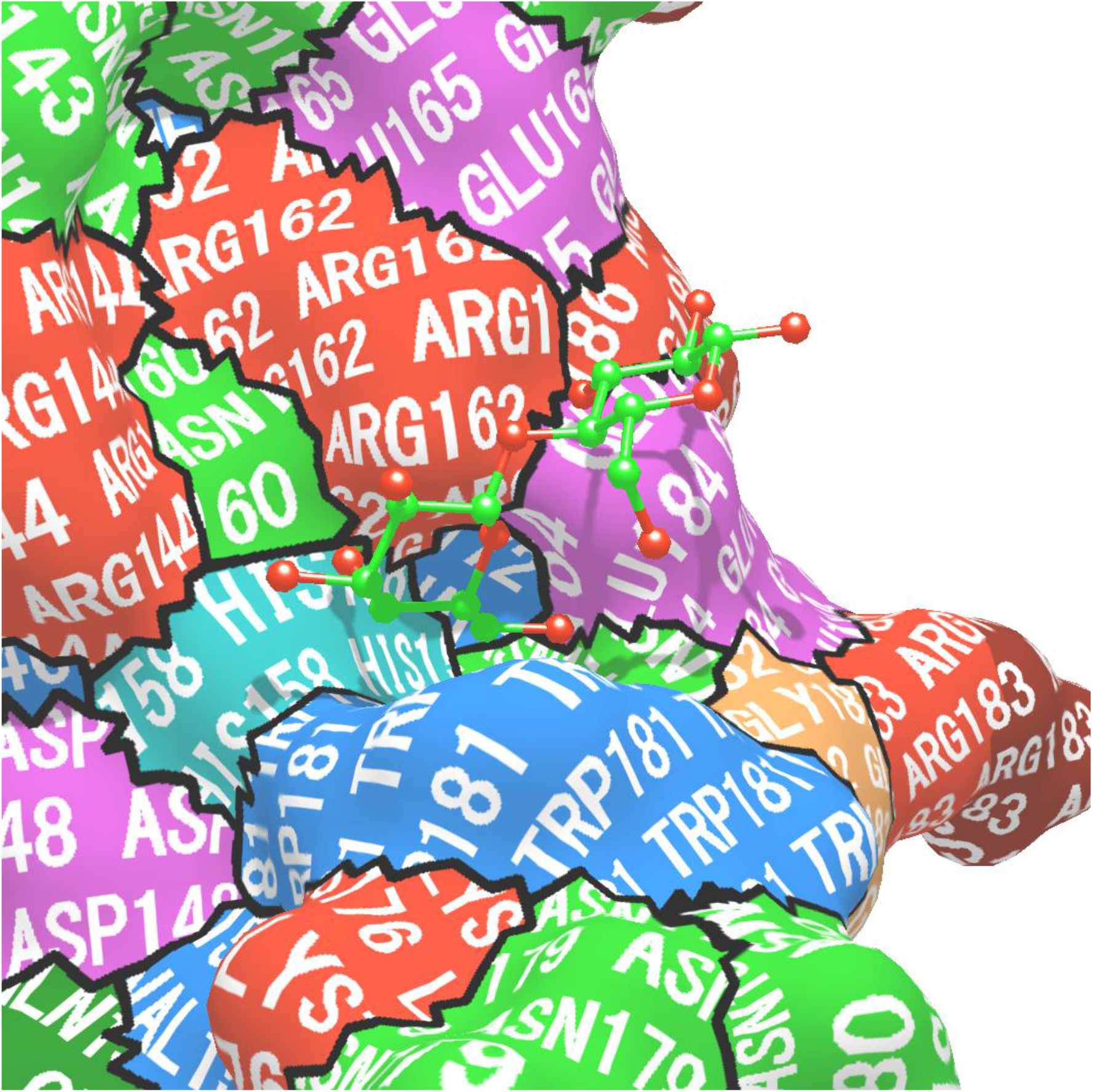
(A) An example of SurfStamp output. SurfStamp was applied to the entry 3ZSJ. The ClustalX (Thompson, et al., 2002) coloring scheme was used. Beta-lactose is shown as a ball-and-stick model with green carbon and red oxygen atoms. The rendered image was created using an in-house .obj viewer, which uses 3D graphics library Three.js (http://threejs.org/). (B) Focused image of the 3ZSJ beta-lactose binding site. The structure was created with options; -tile -font_size 24 -nosep, which would be suitable for 2D images in manuscripts.

## Running, copying, and distributing

I coded SurfStamp entirely in Java language and did not use any external libraries. As a result, it can be run in any operating system (OS) as long as Java VM is installed. In addition, I prepared a graphical user interface as a PyMOL plugin (https://github.com/yamule/SurfStamp-pymol-plugin), which would be useful when users need interactive approaches (e. g. investigating multiple poses of chemical compounds placed in active site).

Furthermore, because I provide this software under the open-source Apache License version 2.0 (http://www.apache.org/licenses/LICENSE-2.0), anyone can use, modify, and distribute this software freely as long as they follow the license term.

## Output example

Figure 1 shows a SurfStamp output example applied against the Galectin carbohydrate recognition domain (CRD), 3ZSJ (Saraboji, et al., 2012) molecule. As you can see, the surface consisting of residues, TRP181, GLU184, HIS158, and so forth, are interacting with the beta-lactose which is shown in the ball-and-stick model.

Figure 2 shows an example of mono-color 3D printed model using “-extrude” option with a 3D object made by EDTSurf. Because the residue information is being extruded, users can recognize which residues are contributing the surface regions of the 3D printed model. It’s also a good idea to color the areas of concern (Figure 2B).

**Figure 2.**
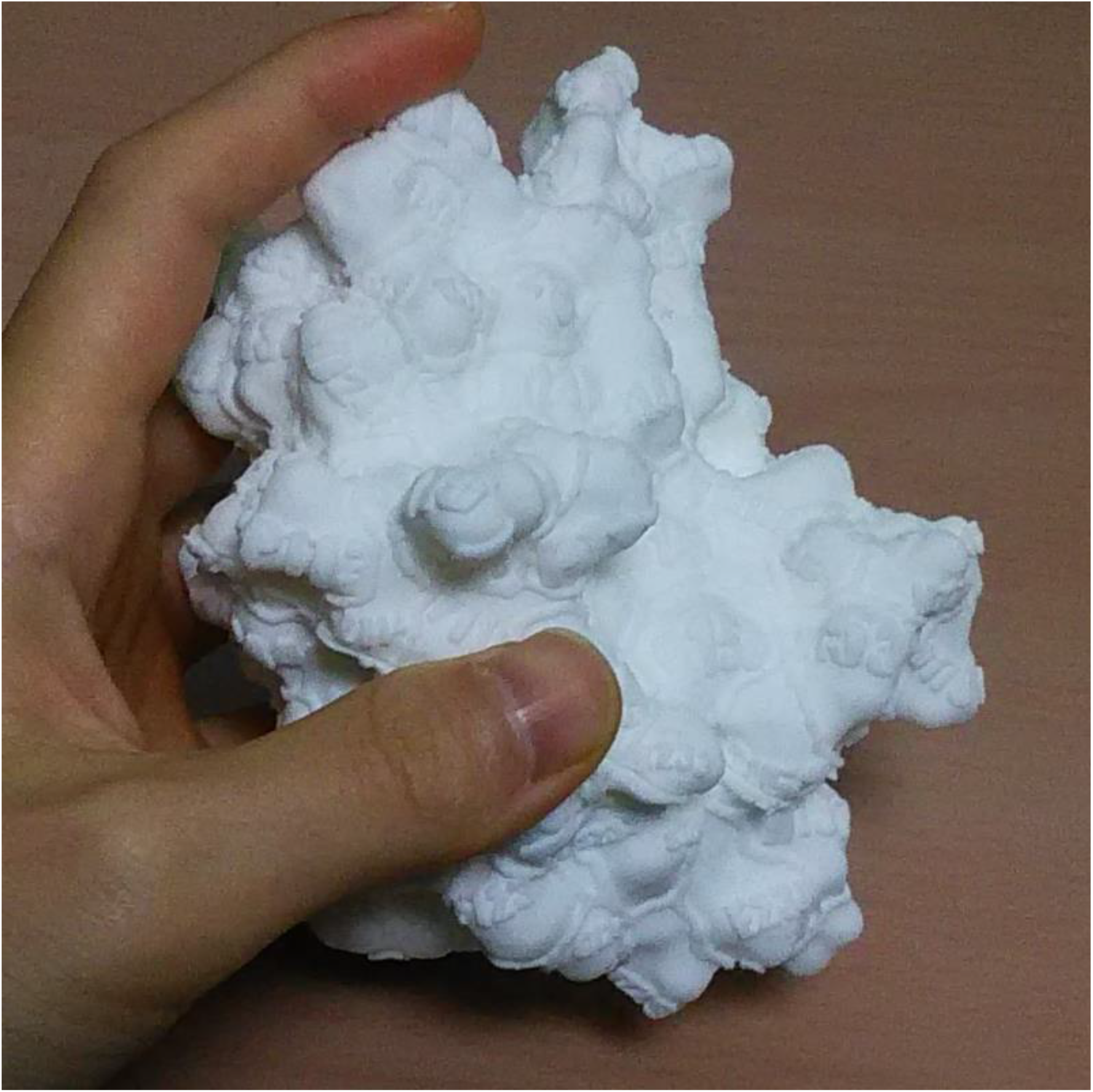
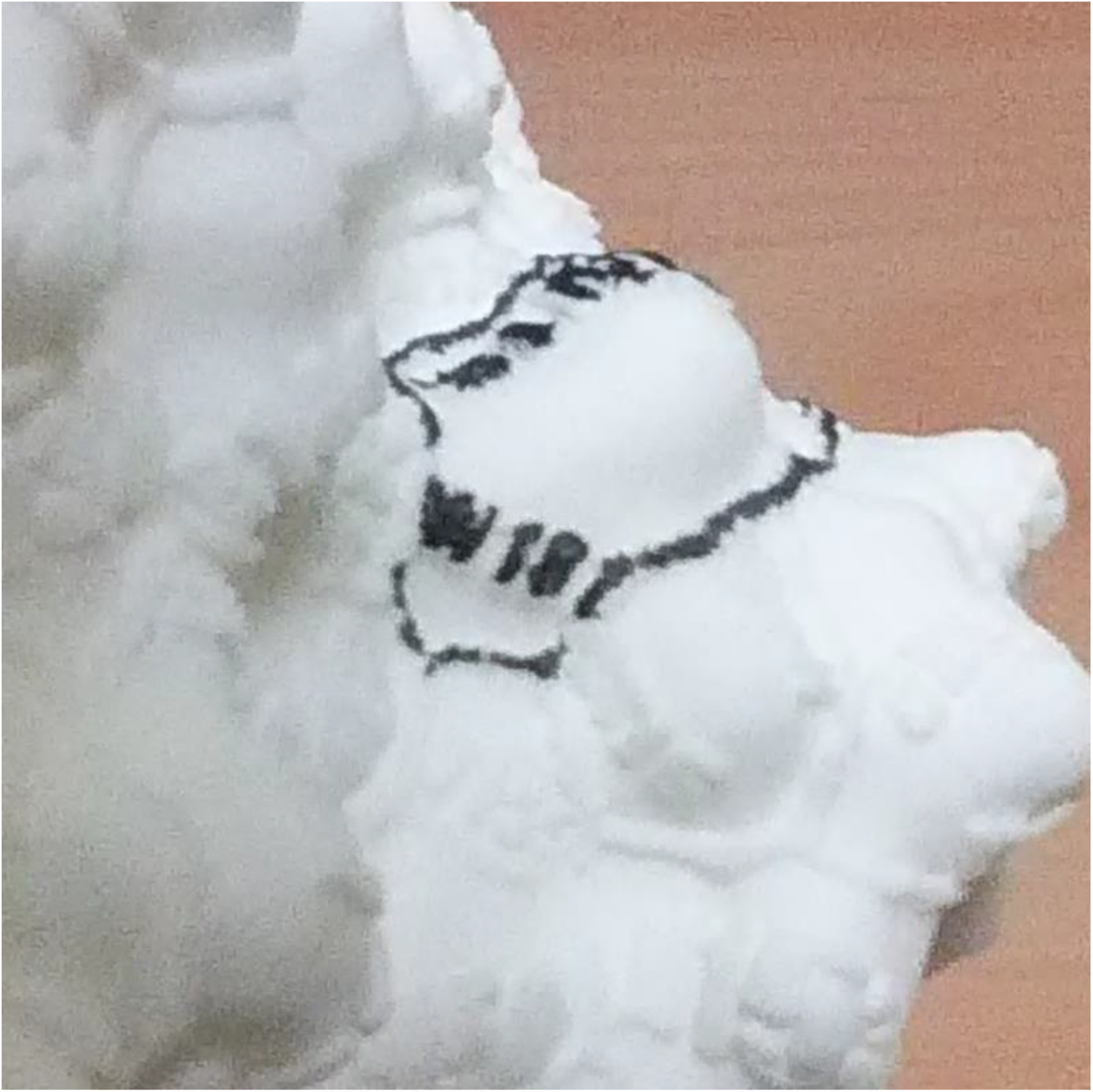
(A) An example 3D printed model created with the “-extrude” option. The 3D printing was performed by DMM make (https://make.dmm.com/) service. 1Å was enlarged to 2.5 mm. The data is available from https://3dprint.nih.gov/discover/3dpx-004144. (B) Focused photo of the 3ZSJ betalactose binding site. I painted the text “W181” and the outline of the region of the surface near the atoms of W(TRP)181.

## Conclusion

Since SurfStamp provides intuitive representations of molecular surfaces, the contents made with this application will be useful in usual research, education, creating figures in academic papers, presentations, and so forth. In addition, since a generous license policy has been adopted, and since the application lacks significant platform dependencies, it can be bundled with other software and can be included in web services. As a result, SurfStamp can be expected to increase the power of molecular visualization techniques, and thereby facilitate research and education.

## Acknowledgements

I really thank staffs of NIH 3D print Exchange (https://3dprint.nih.gov/), especially James Tyrwhitt- Drake (In 2016. As I’ve lost his/her contact information, I don’t know about the prefix.), for helpful discussions and suggestions. The “-extrude” option for mono-color 3D printing was implemented following their suggestions. I sincerely sorry for having provided programs with a lot of bugs in the early days. I thank Dr. Yoshinori Fukasawa. The options; “-color_scheme 4” and “-color_scheme 5” which utilize B-factor and occupancy column, respectively, were implemented following his suggestions. I thank Mr. Masaki Kato. The “-color_dx” option which utilizes result of apbs(Baker, et al., 2001) was implemented following his suggestions.

## Conflict of Interest

The author is an employee of Lifematics Inc. This work was privately done by the author using his own resources.

## Notes

https://github.com/yamule/SurfStamp-public

https://github.com/yamule/SurfStamp-pymol-plugin

### Summary of Updates

Fixed broken references, rearranged some text, added a fig, and added information about the PyMOL plugin.

